# Nutrient-dependent hippocampus dopamine signaling enhances meal-related episodic memory and reduces food intake

**DOI:** 10.64898/2026.06.26.734911

**Authors:** Alexander G. Bashaw, Léa Décarie-Spain, Jessica J. Rea, Logan Tierno-Lauer, Alicia Kao, Olivia Moody, Ryan Wisniewski, Hailey Park, Scott E. Kanoski

## Abstract

**Background:** Dopamine (DA) is a neurotransmitter critically involved in food-related reinforcement learning. While mesolimbic DA reward-associated signaling in the nucleus accumbens has been widely investigated, far less is known about DA function in the hippocampus (HPC), a brain region traditionally known for its role in episodic and spatial memory processes that has recently been associated with appetite and food intake control.

**Methods:** Here we investigated dorsal HPC DA signaling dynamics in rats using fiber photometry to detect changes in DA binding (via GRAB-DA sensors) before, during, and after a meal consumption in food-restricted rats. Pharmacological studies targeting HPC dopamine 2 receptors (D2R) assessed the functional role of HPC DA signaling in food intake and meal-related memory processes.

**Results:** HPC DA binding was significantly elevated in the post-meal relative to the pre-meal state following standard chow consumption. This effect was replicated after consuming a high fat diet or liquid sucrose, but not a low-calorie sweetener. These post-meal DA signaling elevations are dependent on nutrient consumption, as HPC DA binding levels were unaffected by intraperitoneal administration of glucose or the satiation hormone, cholecystokinin, in otherwise fasted rats. Direct HPC D2R agonists administration reduced food intake, whereas HPC D2R blockade after a meal reduced the latency to the next meal and impaired spatial memory for meal location without affecting spatial memory for object location.

**Conclusions:** Collective results identify HPC DA-D2R signaling as a candidate neurobiological mechanism through which nutrient consumption promotes meal-related episodic memory formation, and by extension, reduces subsequent food intake.

## INTRODUCTION

The overwhelming global prevalence of obesity represents not only a major health crisis, but an economic one as well(1). While a new class of weight loss drugs targeting the glucagon-like peptide-1 receptor (either as mono or dual target therapy) are effective and have quickly gained popularity, their use is associated with early weight loss plateau, concomitant gastrointestinal side effects (e.g., nausea and emesis), and the associated weight loss is not sustained when the medication is stopped(2–4). Thus, further research on the neurobiological mechanisms driving the excessive caloric intake associated with obesity is necessary to develop new innovative therapeutic approaches for prevention and treatment.

One hypothesis for the etiology of obesity centers around defects in dopamine (DA) signaling resulting in an improperly calibrated reward-based learning system(5). In human participants, DA signaling in the dorsal striatum is correlated with meal pleasantness ratings(6). Additionally, relative to lean individuals, human participants with obesity have blunted striatal DA levels after eating to satiation and following presentation of images of palatable food(7). This DA disruption hypothesis is further supported by findings that markers of presynaptic DA release mechanisms and postsynaptic responsivity in brain reward centers are reduced in diet-induced obese models in rodents(8–10). Taken together, these results highlight dysregulated striatal DA signaling in obesity, which may be causally related to caloric overconsumption.

Recent work has identified a role of the HPC in the higher-order regulation of food intake, in part via encoding of meal-related episodic memories(11, 12). For example, amnesic patients with bilateral hippocampal damage will eat multiple meals in quick succession yet report little to no change in satiety(13). Additionally, priming healthy human participants to engage in verbal recall of a recent meal will reduce consumption of a subsequent snack(14). However, the neurochemical mechanisms via which the hippocampus encodes meal-related memories to influence food intake are poorly understood. DA signaling is a candidate system given that DA is strongly associated with food-based reinforcement learning and the HPC is innervated by both canonical reward-associated DA neurons in the ventral tegmental area, as well as locus coeruleus neurons that are purported to co-release DA with norepinephrine(15, 16). HPC neurons also express both major classes of DA receptors (i.e., D1R and D2R)(16–19), and HPC DA signaling (detected via PET) is elevated in the HPC of human participants after they consume a palatable milkshake(20). Further, HPC DA 2 receptor (D2R) expression in mice is upregulated in a food-associated context, and chemogenetic inhibition or activation of these D2R-expressing neurons bidirectionally affects food intake(21). While these latter findings suggest a possible role for HPC D2Rs in food intake regulation, modulations of the neural populations that express D2Rs does not directly evaluate the roles of either DA or D2Rs in eating behaviors. Further, while nutrient consumption appears to elevate DA signaling in the HPC, human PET studies lack the temporal resolution to understand the precise endogenous release patterns of DA signaling in the HPC during food consumption.

We hypothesize that food consumption drives increased DA release in the dorsal HPC (HPCd) dentate gyrus whereby signaling via D2Rs promote food-related memory, and by extension, decreases food intake. To investigate the mechanisms of endogenous DA signaling during eating, we performed *in vivo* fiber photometry using molecular DA signaling sensors(22, 23). To study the direct role of D2 receptor signaling, we performed HPCd-targeted pharmacological manipulation at key behavioral timepoints related to meal consumption to parse the system’s involvement in food intake and associated memory formation. Our results collectively reveal that nutrient-induced endogenous HPCd DA signaling acts via D2Rs to influence eating patterns and caloric intake via regulation of episodic meal-related memory formation.

## METHODS AND MATERIALS

### Animals

Adult male Sprague–Dawley rats (Envigo, Indianapolis, IN; PND >60; 320–450 g at the start of experiments) were individually housed in a temperature-controlled vivarium with ad libitum access (except where noted) to water and food (5001 formula, LabDiet, St. Louis, MO) on a 12h:12h reverse light/dark cycle (lights off at 1100). All procedures were approved by the Institute of Animal Care and Use Committee at the University of Southern California.

### Experiment 1: Fiber photometric recording of DA binding during eating

Rats received unilateral HPCd injection of GRAB_DA sensor (targeting the dentate gyrus) and fiber optic cannula implantation, and those that exhibited GRAB_DA responses to quinpirole injection (Figure 1A-B) underwent the refeeding after a fast paradigm (surgical and behavioral procedural details in Supplemental Methods). 24-hour fasted rats were connected via fiber optic patch cord to the photometry recording apparatus in a new clean cage in a novel behavioral room and after a 10-minute baseline period were given 30 minutes ad libitum access to either standard chow (n=7), HFD (n=6), an 11% weight by volume sucrose solution (n=8), or a 0.2% saccharine solution (n=7). DA binding was analyzed for 10 minutes before and after the 30-minute food access period, active eating bouts for solid diets were manually timestamped by an experimenter, and caloric consumption was recorded by weighing food and drink containers before and after the test meal. Two additional control experiments were conducted under similar conditions: (1) standard chow refeeding but with animals expressing a mutated sensor that lacks a functional DA binding domain(23), and (2) an experiment where instead of placing food in the same cage, a familiar object from the home cage was placed in the cage (enrichment tunnel).

**Figure 1:**
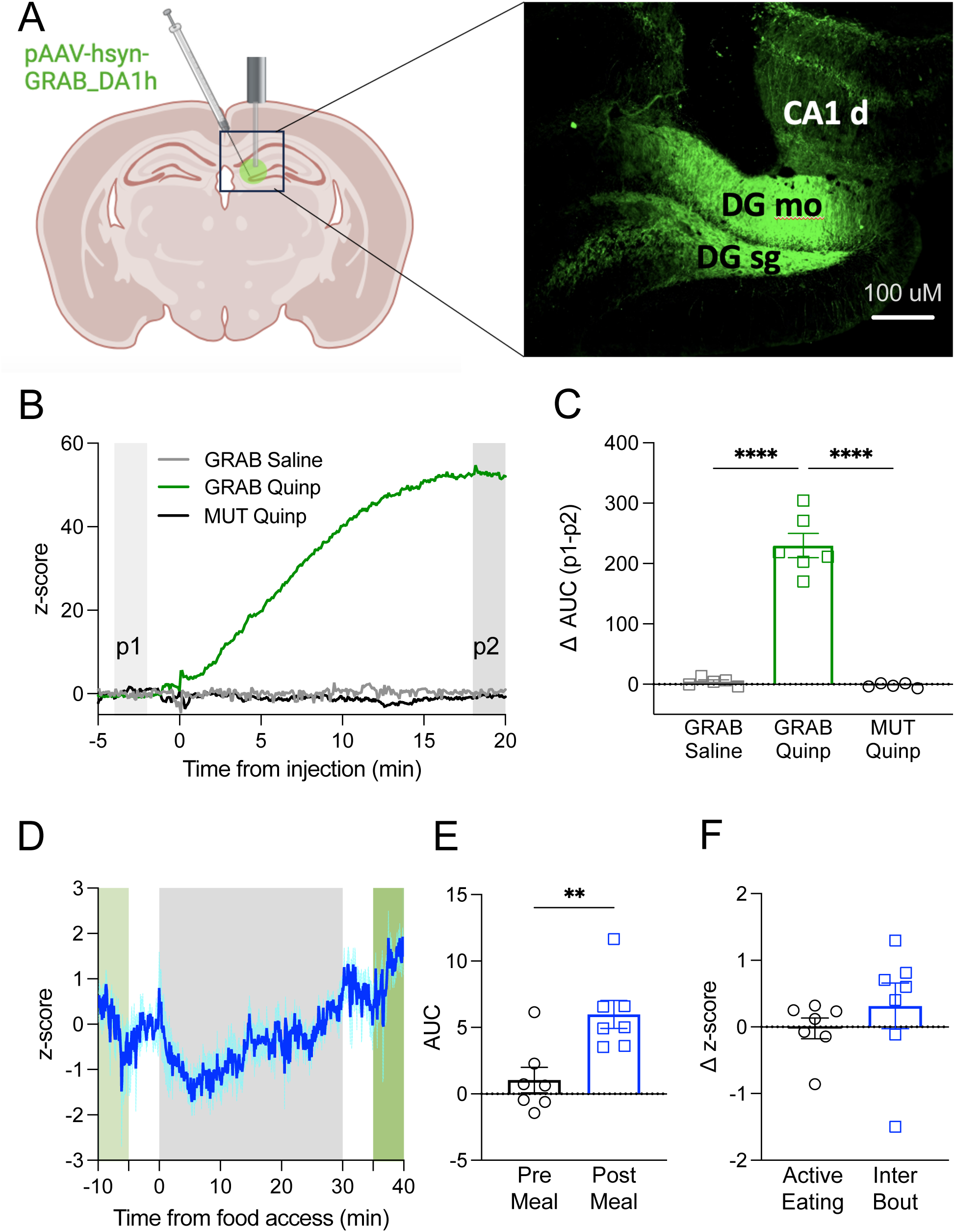
HPCd dopamine binding elevates following meal consumption. **(A)** Surgical prep schematic and representative photomicrograph to show fiber location in HPCd DG subregion and GRAB-DA sensor expression at this site. **(B)** Representative traces display z-score of fluorescence responses of GRAB-DA and GRAB-DA-mutant sensor expressing rats after IP injection of vehicle (0.9% saline) or D2R agonist (quinpirole, 0.5mg/kg). **(C)** Fluorescence changes were measured by the change in area under the curve of the z-scores (AUC) from 4-2 minutes prior to injection (p1) and 18-20 minutes after the injection (p2). Only quinpirole injection induced a significant increase in fluorescence in GRAB_DA expressing rats (n=6 GRAB-Quinp, n=5 GRAB-Saline, n=5 MUT-Quinp, one-way ANOVA, Treatment F(2,13)=104.4, p=0.0459, Tukey’s multiple comparisons test p<0.0001 GRAB-Saline vs GRAB-Quinp, p<0.0001 MUT-Quinp vs GRAB-Quinp). **(D)** Average trace showing z-score of DA binding for rats that were fed standard chow after a 24 hour fast with food access period shaded grey, and pre and post meal periods shaded green. **(E)** Quantification of DA binding during the pre- and post-meal period (AUC) revealed that DA binding significantly increases from the pre-meal to the post-meal state (t(7)=3.853, p=0.0084). **(F)** DA binding as measured by z-scores did not differ during active eating bouts and inter-bout intervals t(7)=0.8081, p= 0.4499). (All between-subjects design for binding validation, all within-subjects design for DA binding during feeding. Data are means ± SEM; **p<0.01, **** p<0.0001.)

### Experiment 2: The effects of systemic injections of glucose or cholecystokinin (CCK) on HPCd DA binding

Rats that received the surgery preparation described in Experiment 1 were fasted for 24 hours, and following a 10-minute baseline period were injected intraperitoneally with (0.5g/ml, 2ml/kg) glucose (n=6), or 3ug/kg CCK (Bachem, n=7) counterbalanced with vehicle 0.9% saline treatment (n=7). DA binding was recorded for 40 minutes after the injection to match the timing of refeeding experiments. For the glucose experiment, blood glucose was measured prior to injection and validated for increases 30, 60, and 90 minutes after injection.

### Experiment 3: Effects of D2 receptor agonism and antagonism on spontaneous eating patterns

Rats with bilateral HPCd cannulae (n=10) had food access removed 1 hour before lights off. For each pharmacology experiment, rats received two counterbalanced treatment days (3 days between treatments) with infusions of drug or vehicle (aCSF) 30-60min before lights off (1100) (Figure 3A) using a within-subject design. For the D2 agonist experiment, rats received 3ug quinpirole (Sigma), and for the D2 antagonist experiment, 5ug raclopride (Sigma). After infusions, rats were returned to home cages in the BioDAQ automated food intake-monitoring system and given access to standard chow, ad libitum. Meal patterns (average meal size and meal frequency) were recorded for 4 hours after lights off.

### Experiment 4: Effects of D2 receptor agonism and antagonism following a salient meal on home cage meal patterns

To model a salient meal (see full procedural details in Supplemental Methods), rats with the surgical preparation described in experiment 3 were maintained on standard chow and were fasted for 24 hours and then (1100) presented with a high-fat diet (HFD; 45% kcal from fat, Research Diets D12451) in a novel environment for 30 minutes (Figure 3B) after lights off. For each pharmacology experiment, rats received two counterbalanced treatment days (3 days between treatments) with infusions of drug or vehicle (aCSF) 5 minutes after the salient meal. After infusions, rats were returned to home cages in the BioDAQ and given access to standard chow. For the D2 agonist experiment, rats received quinpirole (n=9), and for the D2 antagonist experiment, rats received raclopride infusions (n=12), each counterbalanced with vehicle treatment. Meal patterns (average meal size, meal frequency, and first meal latency) were recorded for 4 hours after lights off.

### Experiment 5: Effects of D2R antagonism on Meal Placement Recognition memory (MPR)

Rats with the surgical preparation described in experiments 3 and 4 (n=7) were habituated for 10 minutes on day 1 to the MPR enclosure with empty food cups before returning to their home cage, and baseline empty cup investigation time was scored by an experimenter blinded to subsequent experimental treatments (see full procedural details in Supplemental Methods). Following a 24 hour fast, on day 3, rats were returned to the enclosure where mashed up HFD was placed in the cup least preferred at baseline for 5 minutes. After eating, rats received HPCd infusions of aCSF or raclopride, before returning to home cage. Rats began fasting again on day 4, and on day 5, 24hr fasted rats returned to the enclosure with both food cups empty. Probe empty cup investigation time was manually scored by an experimenter blinded to experimental treatments. MPR experiments (within-subject design) were counterbalanced with two weeks between treatments 1 and 2.

### Experiment 6: Effects of D2R antagonism on Novel Location Recognition memory (NLR)

A modified NLR task was performed to match the timing parameters for MPR to study HPC D2R in a spatial memory task without food intake. Rats with the surgical preparation described in experiments 3-5 (n=6) were habituated for 10 minutes on day 1 to the empty NLR enclosure before returning to their home cage (see full procedural details in Supplemental Methods). On day 3, rats returned to the NLR enclosure with two objects for 15 minutes, and baseline object investigation time was scored by a blinded experimenter. After this training, the rats received infusions of aCSF or raclopride (between-subject design) before returning to the home cage. On day 5, the rats returned to the NLR enclosure with one object moved to a different quadrant. Object investigation time was scored by an experimenter blinded to experimental treatments.

## RESULTS

### HPCd dopamine binding elevates following meal consumption

GRAB_DA sensors injected into the dorsal dentate gyrus with fiber optic cannula implantation allowed us to track changes in DA binding during refeeding after a fast (Figure 1A). HPCd DG DA binding was reliably elevated following 0.5mg/kg IP injections of D2R agonist quinpirole, but not saline, in animals expressing a functional GRAB sensor. However, these changes were not present in animals expressing a mutated sensor that lacks a functional DA binding domain (Figure 1B-C). Rats with validated signal were then refed standard chow following a 24-hour fast to assess HPCd DA binding patterns before, during, and after meal consumption. Given recent findings revealing that acetylcholine release in the dHPC is dynamically modulated by active eating vs. inter bout intervals(24), as is calcium-dependent activity in the ventral hippocampus(25), we investigated if HPCd DA binding also exhibits eating bout related patterns. However, unlike these other hippocampal signaling mechanisms, HPCd DA binding did not differ when comparing active eating bouts to inter bout intervals (Figure 1F). To determine if DA binding changes between hungry and satiated states, we compared DA binding levels from immediately before and after consuming a meal of standard chow. Results revealed that DA binding was elevated from the pre-meal to post-meal state (Figure 1D), and this effect did not correlate with the amount of food consumed (Supplemental Figure 1A). To confirm that this change in fluorescence was not an artifact of the recording setup, we validated that this effect was absent when rats expressing GRAB sensors with a mutated DA binding pocket were refed chow (Supplemental Figure 1B). Given that chow pellets are familiar objects normally present in the home cage, to verify that this effect was based on meal consumption vs. the placement of a familiar object, we confirmed that this temporal change in DA binding was absent when an enrichment tunnel from the home cage was placed in instead of chow (Supplemental Figure 1C).

To determine whether this pre- to post-meal DA binding elevation was based on caloric intake and/or macronutrient profile, we examined whether similar outcomes emerged when animals were fed either a palatable 45% kcal from fat high fat high sugar solid diet (HFD), a liquid 11% weight-by-volume (w/v) sucrose solution, or a non-caloric liquid 0.2% saccharine solution. Like chow-fed rats, fasted rats refed HFD or sucrose exhibited significant pre- to post-meal elevations of HPCd DA binding (Figure 2A-B), yet consumption of saccharin did not change DA binding (Figure 2C). DA binding magnitude changes did not differ between diets, nor were DA binding changes correlated with caloric intake for HFD, sucrose, or volume of saccharine intake (Supplemental Figure 2). Given the timing of the HPCd DA elevations after a 30-minute meal, we wanted to assess if systemic injection of glucose or CCK – metabolic signals elevated during a meal – were sufficient to induce DA binding changes on a similar timescale in hungry rats. IP Glucose injection reliably increased blood glucose levels by 30 minutes after injection (Supplemental Figure 2), but did not change HPCd DA binding (Figure 2E). IP CCK administration, at a dose that reliably reduces food intake(26, 27), also had no effect on DA binding (Figure 2F). Taken together, these results establish that pre- to post-meal DA binding elevations in the dHPC require caloric intake yet are not food or nutrient-specific.

**Figure 2:**
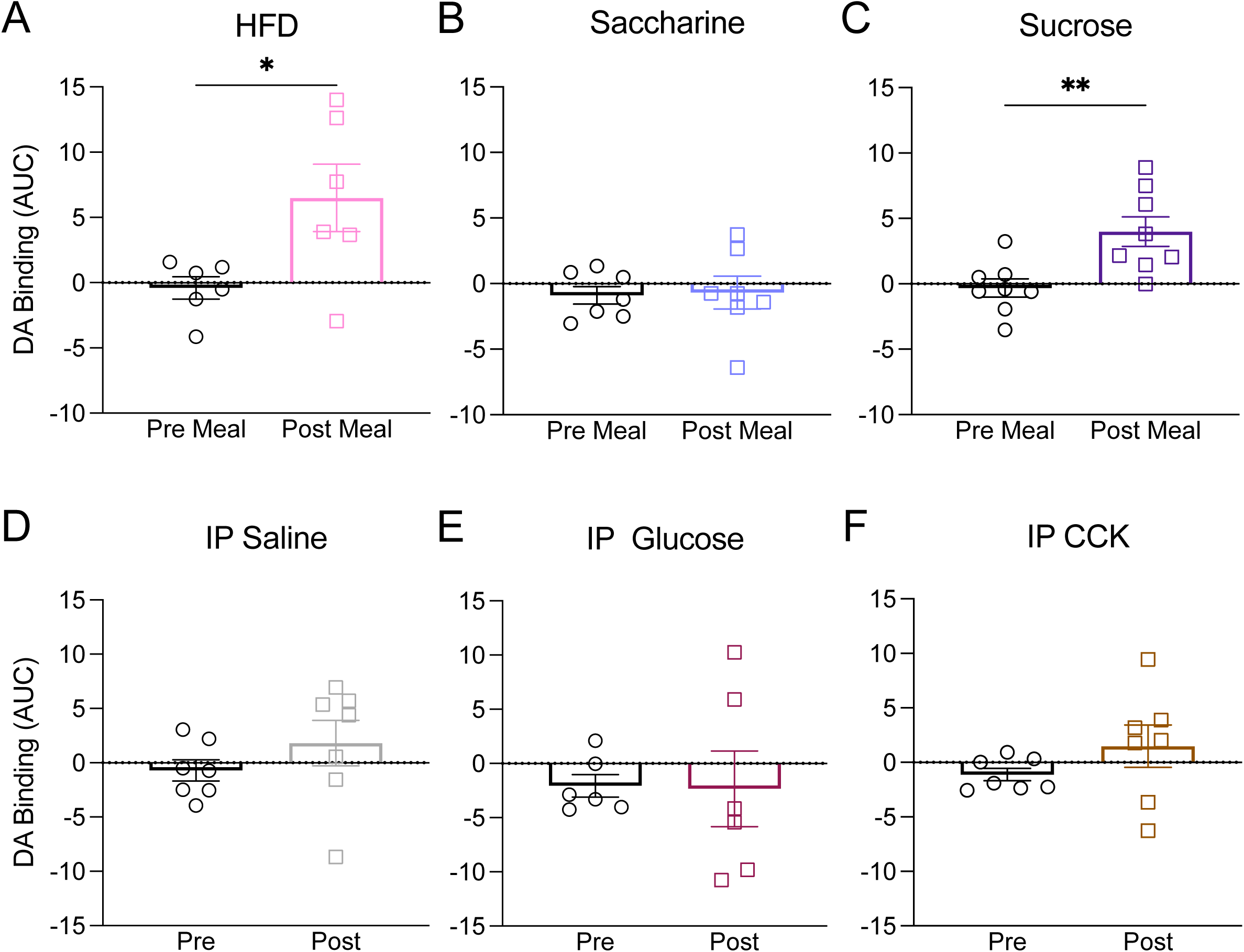
HPCd dopamine binding elevations are calorie dependent. Quantification of DA binding during the pre- and post-meal period (AUC) found that DA binding significantly increases from the pre-meal to the post-meal state for caloric diets (**A**) HFD (t(6)=3.240, p=0.023) and (**C**) liquid 11% sucrose solution (t(8)=3.975, p=0.0054), but does not change for non-caloric (**B**) liquid 0.2% saccharine solution (t(7)=0.2125, p=0.8387). Quantification of DA binding during the pre- and post-injection period (AUC) found that DA binding is not significantly elevated after (**D)** IP 0.9% saline (t(7)=0.2125, p=0.8387), (**E)** IP 0.5g/ml, 2ml/kg glucose (t(6)=0.06436, p=0.9512), or (**F)** IP 3ug/kg CCK (t(7)=1.255, p=0.2560). (All within-subjects design for DA binding during feeding pre-meal vs. post meal and pre-injection vs. post injection. Data are means ± SEM; *p<0.05, **p<0.01).

### Hippocampal dopamine 2 receptors modulate food intake

To determine the functional role of DA signaling in the HPCd on appetite and food intake, we performed direct parenchymal infusions of a DA receptor agonist or antagonist to assess the effects of receptor activation or blockade on spontaneous meal patterns. The D2 receptor agonist quinpirole infused into the HPCd (dentate gyrus) 1hr before dark phase onset significantly reduced cumulative home cage food intake compared to vehicle in unfasted rats at 2 and 4 hours after food access, but the D2 receptor antagonist raclopride did not have an effect under these conditions (Figure 3C-D). To better match the eating conditions described in our refeeding after a fast photometry experiments, we next fasted rats for 24 hours and provided them with 30-mins of HFD access in a novel environment to model a salient meal (Figure 3B). Similar to spontaneous dark phase onset food intake, post salient meal infusion of quinpirole reduced subsequent cumulative home cage intake at 2 hours (Figure 3E), an effect driven by significant reductions in average meal size at this time point (Supplemental Figure 3G). Latency to the first home cage meal initiation after the salient meal was not affected (Figure 3G). On the other hand, while raclopride treatment did not significantly affect cumulative intake (Figure 3F) or eating patterns under these conditions (Supplemental Figure 3B,D), it significantly decreased to latency to subsequently start eating in the home cage after the salient meal (Figure 3H). These findings indicate that HPCd D2 receptors are involved in regulating food intake patterns, with activation reducing food intake and blockade decreasing latency to initiate next meal after a salient meal.

**Figure 3:**
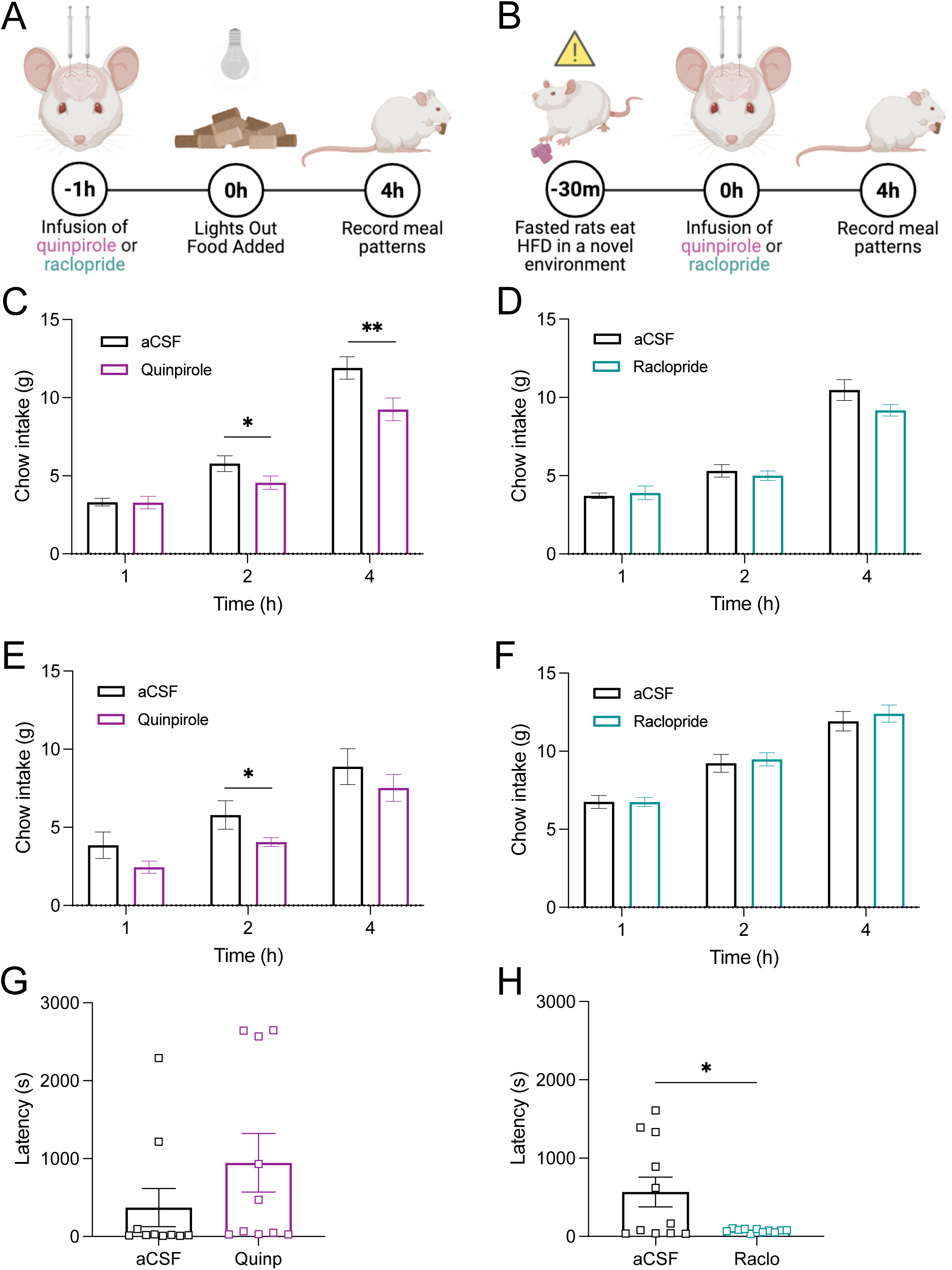
HPCd D2Rs influence food intake patterns. (**A, B**) Schematic of the experimental setup that assessed how HPCd D2R agonism and antagonism affect spontaneous food intake patterns in the BioDAQ under (A) normal conditions or (B) following a salient meal. (**C**) D2R agonism immediately prior to dark cycle significantly reduced cumulative food intake at 2 hours (t(10)=2.973, p=0.0156) and 4 hours (t(10)=3.282, p=0.0095) but not at 1 hour (t(10)=0.9029, p=0.9300). (**D**) D2R antagonism prior to dark cycle onset did not significantly affect cumulative food intake at 1 hour (t(10)=0.3180, p=0.7120), 2 hours (t(10)=0.7163, p=0.4920), or 4 hours (t(10)=2.047, p=0.0709). (**B**) Salient meal experimental setup where fasted rats consumed a 30-minute HFD meal in a novel environment prior to HPCd D2R agonism or antagonism and then returned to the BioDAQ to record food intake patterns. (**E**) D2R agonism after a salient meal significantly reduced cumulative food intake at 2 hours (t(9)=2.604, p=0.0314) but not at 1 hour (t(9)=2.188, p=0.0601) or 4 hours (t(9)=2.224, p=0.0568). (**F**) D2R antagonism after a salient meal did not affect cumulative food intake at 1 hour (t(12)=0.006203, p=0.9952), 2 hours (t(12)=0.4623, p=0.6528) or 4 hours (t(12)=0.9908, p=0.3431). D2R agonism did not effect latency to the next meal (**G**) (t(10)=1.111, p=0.2956), but D2R antagonism significantly reduced latency to eat again after the salient meal (**H**) (t(11)=2.518, p=0.0305). (All within-subjects design for drug vs vehicle. Data are means ± SEM; *p<0.05, **p<0.01).

### Hippocampal D2 receptors encode meal place recognition memory (MPR)

Having established that HPCd DA increases following caloric intake, and that direct manipulation of HPCd D2 receptors is sufficient to change eating patterns, we hypothesized that HPCd signaling may be implicated in the consolidation of meal-related episodic memories. To test this, we performed a meal place recognition assay (MPR)(21), a behavioral paradigm that assesses rats’ ability to remember the location of a recently consumed meal (Figure 4A). Vehicle-treated rats preferentially investigate the previously food-paired cup on the probe day (Figure 4B-C). However, rats that were treated with raclopride (D2R antagonist) following food access exhibited impairments in the test day investigation time of the food-paired cup. Raclopride treatment prevented the increased exploration time for the food-paired cup, and unlike vehicle controls, these animals did not exhibit a positive shift from baseline discrimination (Figure 4B-C). To confirm that D2R antagonism was specifically disrupting meal-related memory rather than general hippocampal function, we performed an analogous hippocampal-dependent memory assay in unfasted rats using a modified novel location recognition (NLR) task to temporally match the parameters of MPR (Figure 4D). Raclopride treatment had no effect on investigation times of relocated objects, and both vehicle and raclopride treatments induced a significant positive shift in investigation of the moved object (Figure 4E-F). Follow-up experiments also confirmed that raclopride did not impair non-food-related short-term hippocampal-dependent spatial memory using more common NLR parameters (Supplemental Figure 4). These findings together emphasize the role of HPCd D2Rs in responding to meal related DA elevations as an essential signal for meal-related episodic memory consolidation.

**Figure 4:**
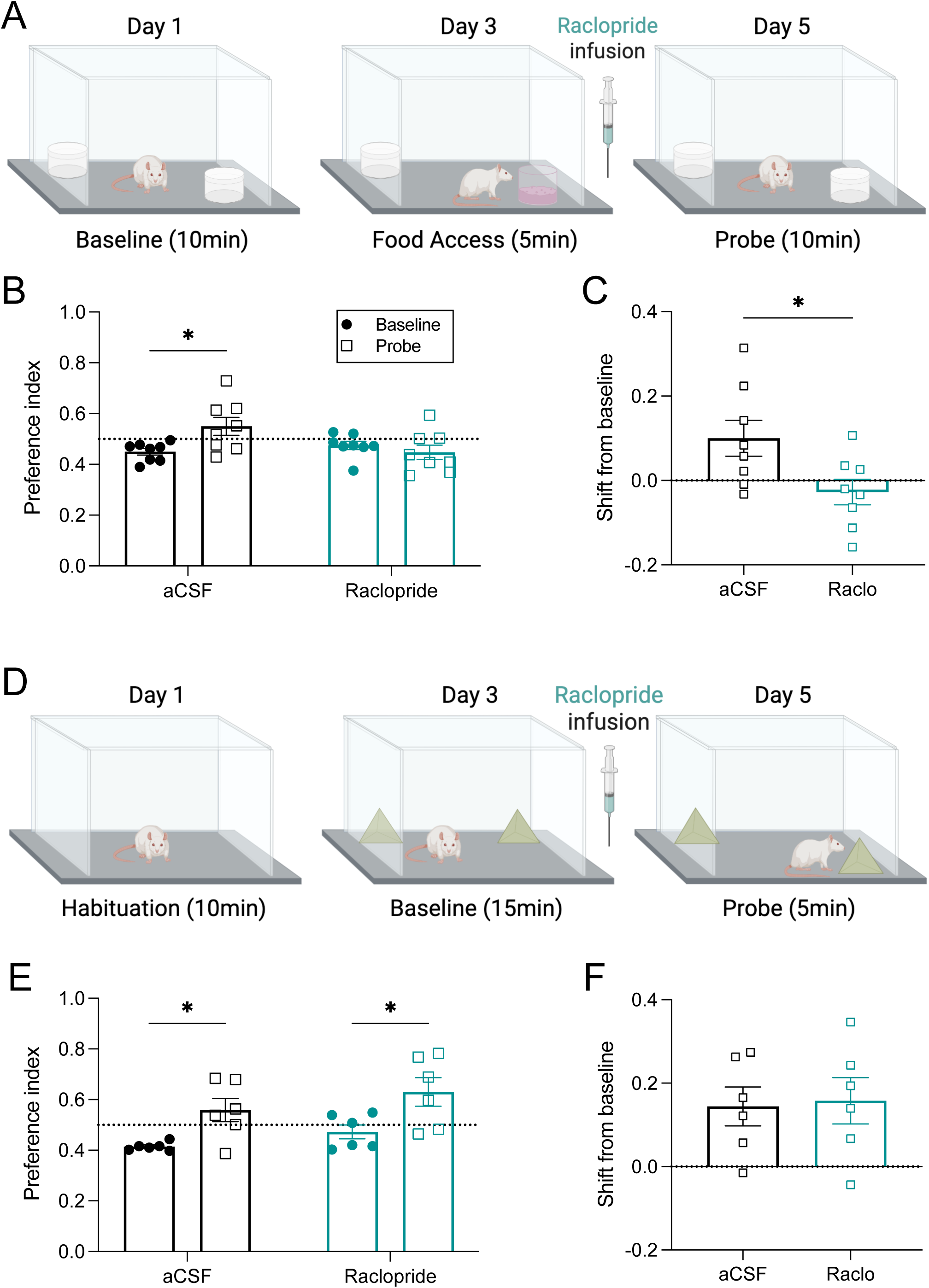
Hippocampal D2 receptors encode meal place recognition memory (MPR). (**A**) Schematic of the experimental setup that assessed how HPCd D2R antagonism affected MPR. (**B**) Vehicle treated rats exhibited significant increase in time spent investigating the food-paired cup at probe vs. baseline, indicating they had learned the meal location (t(8)=2.368, p=0.0498). However, raclopride treated rats did not show a difference in their preference for the food-paired cup from baseline to probe, indicating a memory impairment (t(8)=0.9158, p=0.3903). Raclopride treatment impaired the positive shift in investigation time from baseline associated with meal place recognition memory ((t(8)=2.893, p=0.0232). To confirm that this impairment was specific to food-related memory, we performed a hippocampal-dependent novel location recognition (NLR) experiment that was temporally similar to the MPR experiment but did not involve food (**D**). Both vehicle and D2R antagonist treated rats significantly increased their preference for the relocated object from baseline to probe (**E**) aCSF t(6)=3.106, p=0.0267), raclopride t(6)=2.829, p=0.0367), and both groups had a positive shift from baseline (**F)** t(6)=0.1850, p=0.8569). (MPR experiments within-subjects design for drug vs vehicle. NLR experiments between subject design for drug vs. vehicle. Data are means ± SEM; *p<0.05, **p<0.01).

## DISCUSSION

Brain dopamine (DA) signaling is associated with reward-based learning and motivation in both animal models and human subjects(9, 10, 28–30). Midbrain ventral tegmental area (VTA) DA neurons densely innervate the striatum, and fire in response to both primary food rewards at intake onset(31) and post-oral gastric administration of nutrients(32). However, in addition to canonical mesolimbic reward pathways, VTA DA neurons also project to other brain regions including the amygdala, prefrontal cortex, and hippocampus (HPC). HPC DA binding in human subjects consuming a milkshake is elevated 35-40 minutes after milkshake consumption as measured by PET radioactive raclopride (competitive D2R antagonist) displacement by endogenous DA(20). Using GRAB_DA (D2R based) sensors to track DA with greater temporal resolution in rats, here we also found HPC DA binding elevations at the timescale of a post-ingestive/post-meal response. DA binding was elevated when fasted rats consumed standard chow, high fat diet, or a liquid sucrose solution. However, we did not observe DA binding changes when rats consumed a non-caloric artificially sweetened taste solution, implying that caloric intake was required for post-consumption HPC DA elevations.

To investigate the functional behavioral significance of post-prandial DA binding, we performed direct pharmacological manipulations of HPC D2Rs in close temporal proximity to eating events. D2R agonism before meals reduced spontaneous food intake via meal size reduction in non-fasted rats as well as after fasted rats ate a salient meal. These results are consistent with findings that chemogenetic inactivation of D2R expressing neurons reduced food intake in mice(21), yet importantly, expand these findings by identifying both dopamine, and D2R signaling as crucial components of HPC-mediated food intake reduction.

D2R antagonism (raclopride) did not affect food intake in home cage spontaneous meal pattern analyses, or when home cage spontaneous meal pattern analyses were conducted after a large and salient meal. However, HPC D2R antagonism following salient meal consumption did significantly reduce the latency to subsequently initiate a meal in the home cage. This suggests that HPC D2R blockade during the post-prandial window when endogenous HPC DA binding is elevated may impair the episodic memory of that meal, which by extension, leads to earlier subsequent meal initiation. This is consistent with our findings that D2R agonism reduced food intake, and together suggests a role of intact HPC D2R in forming meal-related memories that function to mediate eating patterns.

In addition to influencing the post-prandial meal initiation interval, a period referred to as “satiety”, HPC D2R antagonism impaired memory for the location of where a previous meal was consumed. This effect was specific to food location memory, as D2R antagonism did not impair hippocampal-dependent object location memory when tested under analogous parameters. Given that D2R antagonism both reduced latency to initiate a meal following a large and salient meal (potentially via impaired memory of when the last meal was consumed), and impaired memory retention for a salient meal location, these findings suggest that HPC DA exerts its action on D2Rs in a sensitive post-prandial window to form episodic memories of meals (including when, and where) that regulate subsequent eating patterns, particularly when such meals are salient.

While our study establishes a functional role for DA HPC signaling in mediating food consumption, satiety, and meal-related memory, the sources of these functional HPC DA responses are yet to be empirically established. Although the timing of postprandial HPC DA increases is consistent with a midbrain DA source responding to post-ingestive cues(33, 34), we cannot rule out the involvement of other DA inputs to the HPC in regulating food consumption and food-related memory. For example, the Locus Coeruleus (LC) is a broadly projecting hindbrain region that can co release DA along with norepinephrine(NE)(15, 16, 35), and more richly innervates the dorsal hippocampus compared to the VTA(36). Like the VTA and HPC, the LC plays a role in ingestive behaviors(37) and memory consolidation(35), particularly in the context of novelty detection. Optogenetic activation of LC TH+ neurons (that produce NE and/or DA) in mice improved memory performance for a food location task 24 hours after training(36). Further, LC neurons dynamically increase their activity during food consumption(37), suggesting that LC DA may contribute to increases in HPC DA binding after meals. Future investigation is needed to parse the relative contribution of VTA and LC neurons to HPC DA release related to ingestive behaviors.

In addition to gaps in field regarding the source of HPC DA, downstream mechanisms of DA binding on the synaptic physiology of hippocampal neurons remain elusive. HPC neurons express all classes of DA receptors, including D1 and D2R type receptors(18, 19) and some HPC neurons have been documented to co-express both D1 and D2Rs(38), further complicating how DA signaling might impact hippocampal synaptic physiology. We chose to focus on hippocampal D2Rs based on findings that mouse HPC D2R expression is upregulated in a food context(21), human PET studies report HPC DA elevations using a radioactive D2R antagonist(20), and also because GRAB-DA sensors are D2R based(23). Further, D2Rs have been shown to have relatively high affinity for DA when compared to the D1R subtype(39), which may make them better suited to detecting the relatively low levels of DA release in the HPC as compared to the striatum. Synaptic physiology experiments have shown that genetic D2R knockout, inhibitory D2R RNA treatment, and pharmacological D2R blockade in mice leads to impairments in hippocampal synaptic plasticity and spatial learning(40, 41). Further work is needed to examine how these synaptic perturbations may affect food-related memories and ingestive behaviors.

Collective results support a framework in which HPC D2Rs respond to post prandial (nutrient-induced) DA increases, and that this signaling pathway is critical for food consumption, satiety, and food-related episodic memory. It is important to further investigate how this system is impacted by dietary and metabolic challenges, as perturbations to this system may resemble those observed in the striatum that are associated with the etiology of overweight and obesity(42–44). Given recent findings in humans establishing an orexigenic network in the HPC that is dysregulated in obesity(45), our findings highlight a need for further study of the contribution of HPC DA signaling to food intake control and obesity etiology.

## Acknowledgments

Figures 1, 3, and 4 were created with assistance from BioRender.com. We would also like to acknowledge the following funding sources:

Alexander Bashaw: NIDDK NRSA F31 (1F31DK142526-01A1)

Scott Kanoski: NIDDK (R01DK104897 and RO1DK123423)

## Supplemental Materials & Methods

### Surgery

For all surgical procedures, rats were anesthetized and sedated via intramuscular injections of ketamine (90 mg/kg), xylazine (2.8 mg/kg), and acepromazine (0.72 mg/kg) cocktail. Rats were also given preoperative analgesic (subcutaneous injection of Buprenorphine-XR [1.3 mg/kg]). All rats recovered for at least one-week post-surgery prior to initiation of behavioral procedures.

### Intracranial viral injection and in vivo photometry optic fiber placement

Once sedated, the surgical site was shaved and disinfected with iodine and ethanol swabbing before the animals were secured in a stereotaxic apparatus. For fiber photometry preparations, Adeno-associated virus (AAV) encoding GRAB_DA (500 nL; original titer ≥ 1×10^13^ vg/mL, pAAV.hSyn.GRAB_DA1h, 113050-AAV9, or a Mutant sensor control pAAV.hSyn.GRAB_DA.mut original titer ≥ 1×10¹³ vg/mL,140555-AAV9 Addgene, Watertown, MA, USA)—previously validated as a selective in vivo D2R based sensor in rodents was injected unilaterally into the dorsal dentate gyrus of the hippocampus (HPCd) (A/P: −3.12; M/L: ±1.2; D/V: −3.9, with bregma as the reference point) using a micro infusion pump (Harvard Apparatus, Cambridge, MA, USA) equipped with a 33-gauge micro syringe injector connected to a PE20 catheter and Hamilton syringe. The injection was delivered at a rate of 5 μL/min, with injectors left in place for an additional 2 minutes to ensure complete infusion. Following viral delivery, fiber optic cannulae (flat 400 μm core, 0.48 numerical aperture, 5 mm; Doric Lenses Inc., Quebec, Canada) were implanted above the injection site (A/P: −3.12; M/L: ±1.2; D/V: −3.8, with bregma as the reference point). The optic fibers were secured to the skull using jeweler’s screws, instant adhesive super glue, and dental cement. All rats recovered for at least one-week post-surgery prior to experimental procedures. Fiber locations were verified by post-mortem, and animals with fiber tracks in the dHPC were included in the study.

### Cannula implantation for drug injections

For parenchymal pharmacological DA receptor agonist/antagonist delivery, rats were surgically implanted with indwelling single guide cannula bilaterally (26-gauge, Plastics One, Roanoke, VA) using the following stereotaxic coordinates, which are relative to the location of bregma: −4.08 mm anterior/posterior (AP), +/- 2.50 mm medial/lateral (ML), and −2.60 mm dorsal/ventral (DV). Cannula were affixed to the skull as previously described using jeweler’s screws and dental cement. Drug injections were made with injectors that projected 1.0mm beyond the end of the guide cannula. Experiments involving pharmacological administration included bilateral dHPC (dentate gyrus [DG] region) injections of Raclopride and Quinpirole which were dissolved in artificial cerebral spinal fluid (aCSF) and diluted to 2.5ug/100nl (5ug total) and 1.5ug/100nl (3ug total), respectively. Injections were administered using a microinfusion pump (Harvard Apparatus, Holliston, MA) connected to a 26-gauge microsyringe injector through the indwelling guide cannulae. Flow rate was calibrated and set to 5 ml/min and 100nl injection volume per hemisphere. Injectors were left in place for 30-sec to allow for complete infusion of the drug. Placements for HPCd cannulae were verified post-mortem by injection of 100nl blue dye (100nl, 2% Chicago sky blue ink) through the guide cannulae. Data from animals with dye confined to the HPCd DG were included in the analyses.

### In vivo fiber photometry

Fiber photometry recordings were conducted following previously established methods. Signal acquisition was performed using the Neurophotometrics fiber photometry system (Neurophotometrics, San Diego, CA) at a sampling frequency of 40 Hz, with alternating excitation wavelengths of 470 nm (DA-dependent) and 415 nm (DA-independent). Fluorescence was transmitted through an optical patch cord (Doric Lenses) and directed onto the implanted optic fiber, which in turn relayed neural fluorescence back through the same optic fiber and patch cord to a photoreceiver. Behavioral events, such as food access and removal, were time-stamped using the data acquisition software (Bonsai). To correct for baseline neural activity and motion artifacts, the DA-independent signal was subtracted from the DA-dependent signal, and the resulting fluorescence fluctuations were fitted to a biexponential curve. The corrected fluorescence signal was then normalized for each rat by calculating ΔF/F using the average fluorescence signal from the entire recording, followed by conversion to z-scores.

Finally, the normalized signal was aligned to relevant behavioral events, and data extraction was performed using custom MATLAB code.

### Fiber photometry during meal consumption

To determine if animals had viable HPCd DA signal, animals were recorded for 5 minutes prior to IP injections of 0.9% saline or 0.5 mg/kg Quinpirole, and then for 20 minutes after injection. Rats injected with Quinpirole that exhibited significantly increased AUC of zscore comparing the first 5 minutes to the last 5 were deemed to have signal and included in subsequent experiments. Prior to refeeding the test day, rats underwent a 24-hour fast with *ad libitum* access to water. Testing occurred in the early nocturnal/dark phase for a 24-hour period without food. On test days, animals were first placed in a neutral context (neutral rat cage in a procedure room with dim lighting) for 15 minutes without solid food, followed by a 30-minute solid food access period (Laboratory Rodent Diet 5001, St. Louis, MO, USA). Afterward, food was removed, and animals remained in the context for an additional 10 minutes. To assess the effects of physiological HPCd DA binding during refeeding after a 24-hour fast, in vivo fiber photometry was performed. The 5-min pre-consumption period for each animal was defined as a 5-min period terminating 5 min before food access. The 5-min post-consumption period was defined as the last 5-min period of the recording. Liquid meal consumption analyses were conducted under similar conditions, except that the consumption tests occurred in operant boxes with lickometers to deliver solutions (Med Associates Inc., Fairfax, VT).

### 2.5 Meal Place Recognition memory task

MPR is used to assess food location memory. Rats were habituated to a semi-transparent square box (41.9 cm L × 41.9 cm W × 38.1 cm H), placed in a dimly lit room in which two adjacent desk lamps were pointed toward the floor for 10 mins 1-2 days before the experiment. On baseline day, the rats were introduced to the MPR apparatus which now contained two empty stainless steel espresso cups at opposite corners of the apparatus. The time each rat spent exploring the cups was quantified by hand-scoring of video recordings by an experimenter blinded to the animal group assignments and object exploration was defined as the rat sniffing or touching the object with the nose or forepaws. The rats were returned to their home cages for 24 hours. Prior to the training phase (day 3), rats underwent a 24hr fast with *ad libitum* access to water. The rats were then returned to the apparatus, but now the less preferred cup on baseline day was filled with 5 grams of crushed up HFD which rats were allowed to eat for 5 minutes. After returning to their home cage for 5 minutes, rats HPCd DG were bilaterally infused with aCSF or Raclopride. Approximately 1hr later, rats were given back standard chow in their home cages. Prior to the probe phase (day 5), rats underwent a 24-hour fast with *ad libitum* access to water. The rats were then returned to the MPR apparatus with empty food cups for 10-min and investigation time was recorded.

### Novel Location Recognition tasks

NLR (short term) was performed to assess spatial recognition memory. A grey opaque box (38.1 cm L × 56.5 cm W × 31.8 cm H), placed in a dimly lit room in which two adjacent desk lamps were pointed toward the floor, was used as the NLR apparatus. Rats were habituated to the empty apparatus and conditions for 10 min 2 days prior to testing. On test day, rats were infused with raclopride or aCSF approximately 1 hour before testing. Testing constituted a 5-min familiarization phase during which rats were placed in the center of the apparatus (facing a neutral wall to avoid biasing them toward either object) with two identical objects placed in two corners of the apparatus and allowed to explore. The objects used were either two identical empty glass salt-shakers (that never contained salt) or two identical empty soap dispensers (that were never used with soap; first NLR time point), or two identical textured glass vases or two identical ceramic jugs (second NLR time point). Rats were then removed from the apparatus and returned to their home cage for 5 min. During this period, the apparatus and objects were cleaned with 10% ethanol solution and one of the objects was moved to a different corner location in the apparatus (i.e., the object was moved but not replaced). Rats were then placed in the center of the apparatus again and allowed to explore for 3 min. The types of objects used and the novel location placements were counterbalanced by group. The time each rat spent exploring the objects was quantified by hand-scoring of video recordings by an experimenter blinded to the animal group assignments and object exploration was defined as the rat sniffing or touching the object with the nose or forepaws. NLR (long term) was assessed to match the timing of MPR experiments described previously. The same experimental setup was used as short-term NLR, but rats were given at 15-min familiarization phase, infused with aCSF or raclopride and then returned to their home cage. 48 hours later, the rats were returned to the apparatus for the probe with the relocated object, and allowed to explore for 5 min while an experimenter tracked investigation time for each object.

### Salient Meal Procedure

Prior to refeeding the test day, chow-maintained rats housed in the BIODAQ underwent a 24 hour fast with *ad libitum* access to water. Then, the rats were moved to a clear behavior box in a lowly lit experiment room and given 30 minutes of ad libitum access to HFD. After returning to their home cage for 5 minutes, each rat was bilaterally infused in the HPCd with aCSF, raclopride or quinpirole. The rats were then returned to their home cages in the BioDAQ, BioDAQ food monitoring intake system to which they were habituated to for at least 5 days prior to any treatments. Food access was restored, and food intake parameters were recorded for the subsequent 4hrs.

### Meal patterns in BioDAQ automatic food intake monitoring system

Rats were individually housed in BioDAQ Food and Water intake monitoring system (Research Diets Inc. New Brunswick, NJ) which recorded episodic ad libitum feeding activity of rats in their home cages. All PSCs were validated for accuracy using 10.0g standard weights with an allowable margin of error +/- 0.05g prior to the start of the experiment as well as before test days. All PSCs were required to have QUIET readings prior to the start of any session. BioDAQ data was analyzed using an inter meal interval (IMI) of 900 seconds to separate meals, and bouts were filtered to be between −9.00 and 9.00g in size. Data was also set to have a period that started at the onset of the dark cycle when the behavioral task would begin, in this case 11:00AM to coincide with the light schedule in the room in which they were housed. The number of periods in the day was set to 24 resulting in 1hr long period bins from which cumulative intake as well as average meal size, duration and latency to eat could be calculated. Following exportation to Excel, values were obtained from the PSC by period tab to record individual animals’ intake. First meal latency was taken from the “Meals” tab. For pharmacological experiments on test days, BioDAQ home cage food access was removed 1 hour prior to dark onset. Drug treatments were counterbalanced across animals using a within-subject design. Infusions of aCSF or Raclopride/Quinipirole occurred 45—60 min prior to the start of the dark cycle, at which point access to food was restored and food intake parameters were recorded for the subsequent 4hrs.

### Statistical Analyses

Statistical analyses were performed using GraphPad Prism 10.0 software (GraphPad Software Inc., San Diego, CA, USA) and Microsoft Excel V16.66.1. Data are expressed as mean ± SEM. Statistical details can be found in the figure legends and *n*’s refer to the number of animals for each condition. Differences were considered statistically significant at *P* < 0.05. A 1-way ANOVA with tukey multiple comparisons was used to compare between subject DA binding changes (delta AUC) for GRAB_DA1h 0.9% saline injection, GRAB_DA1h quinpirole 0.5mg/kg injection, and GRAB_DA.mut quinpirole injection. Two-tailed paired Student’s *t*-tests were used to compare within-subject HPCd DA binding for pre-meal and post-meal periods during the consumption of standard chow, HFD, 11% sucrose, 0.2% saccharin as well pre and post injections of 0.9% saline, CCK (3mg/kg), Glucose (0.5g/ml, 2ml/kg). Two-tailed paired Student’s *t*-tests were used to compare within-subject preference index from baseline to probe and shift from baseline to probe for MPR task for both aCSF and raclopride treatements. Two-tailed paired Student’s *t*-tests were used to compare within-subject preference index from baseline to probe and shift from baseline to probe for long term NLR task for both aCSF and raclopride treatements. Two-tailed unpaired Student’s *t*-test was used to compare between-subject short term NLR for aCSF and raclopride treatments. One-sample t-test with theoretical mean of 0.5 was used for NLR investigation preference for aCSF and raclopride treated groups. Two-tailed paired Student’s *t*-tests were used to compare within-subject BIODAQ meal patterns for aCSF and Raclopride or aCSF and Quinpirole. Simple linear regressions were performed to analyze whether magnitude of DA binding changes predicted caloric intake of chow, HFD, and sucrose and volumetric intake of saccharine. Simple linear regressions were performed to analyze whether magnitude of DA binding changes predicted changes in glucose levels. Outliers were identified using the Grubb’s test for outliers post-hoc at significance level of alpha = 0.05. For all experiments, assumptions of normality, homogeneity of variance (HOV), and independence were met where required.

---

**Supplemental Figure 1:**
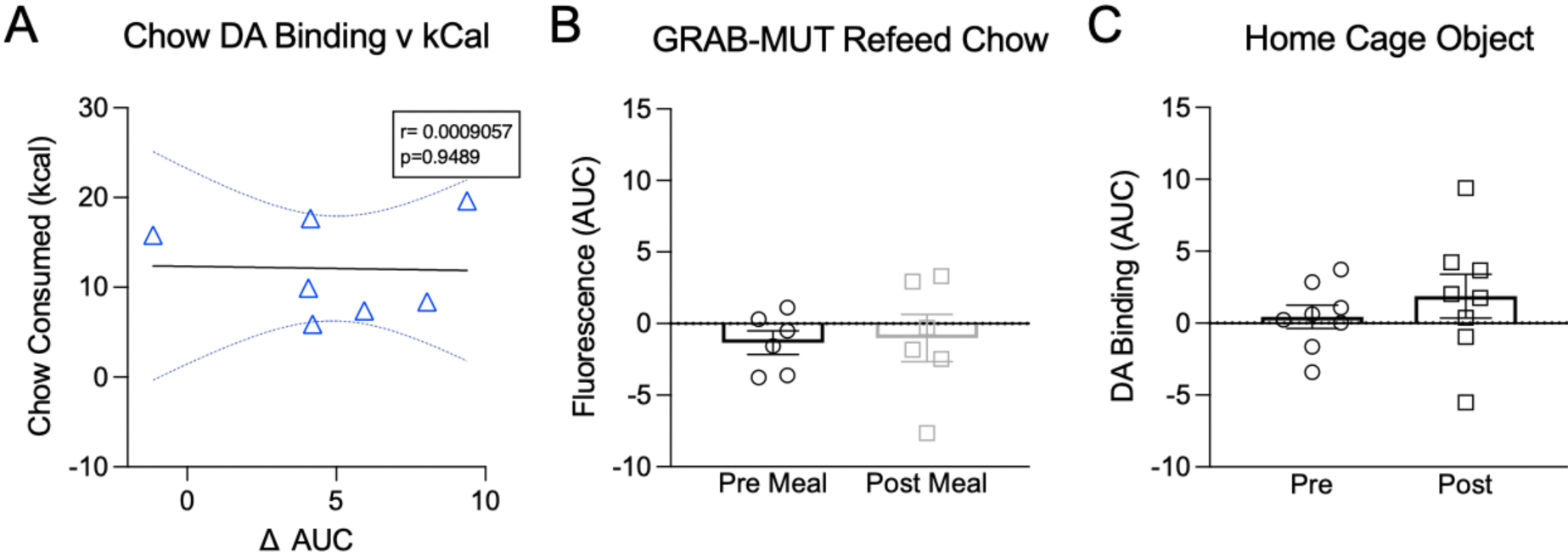
DA binding changes do not predict calories consumed, and post-meal HPC DA binding fluorescence increases are not present with mutant GRAB sensors or without food intake. A) Regression of DA binding changes from the pre-meal to the post-meal on calories consumed of standard chow (n=7, R^2^=0.0009057, p=0.9489). B) GRAB_Mut sensors have a mutated binding DA domain and thus do not bind DA. Rats expressing GRAB_Mut sensor in HPC did not exhibit a pre-meal to post meal increase in fluorescence (t(5)=0.2754, p=0.7940). C) When fasted rats are presented with a home cage object for 30 minutes instead of given food, there was no significant difference in DA binding from the pre to post 30-min object exposure state (t(7)=0.8171, p=0.4408). (All within-subject design. Data are means ± SEM.)

**Supplemental Figure 2:**
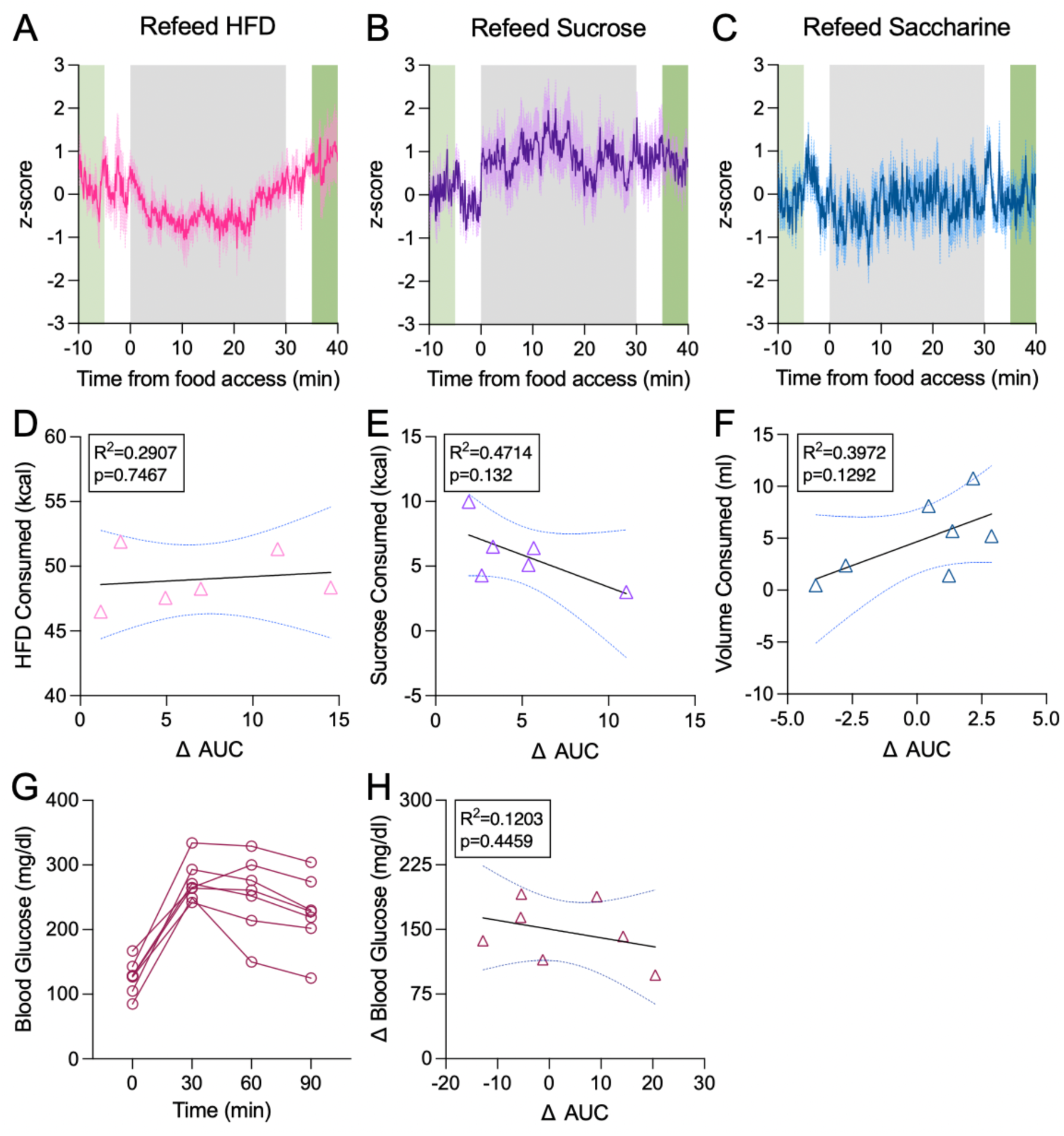
HPC DA binding magnitudes are not correlated with intake amounts or blood glucose levels. **A-C**) Representative traces of DA binding when fasted rats are refed various diets. Green bars represent time windows where z-score values used to compute pre and post meal DA binding, and gray bars indicate diet access period. Regression of DA binding changes on calories consumed for **D)** High Fat Diet (n=6, R^2^=0.02907, p=0.7467), **E)** 11% sucrose solution (n=6, R^2^=0.4714, p=0.132), and on intake volume for **F)** 0.2% saccharine solution (n=7, R^2^=0.3972, p=0.1292) **G**) Blood glucose response curves for rats receiving IP glucose injection. **H)** Regression of DA binding changes on blood glucose changes from before injection vs 30 minutes after injection (n=7, R^2^=0.1203, p=0.4459).

**Supplemental Figure 3:**
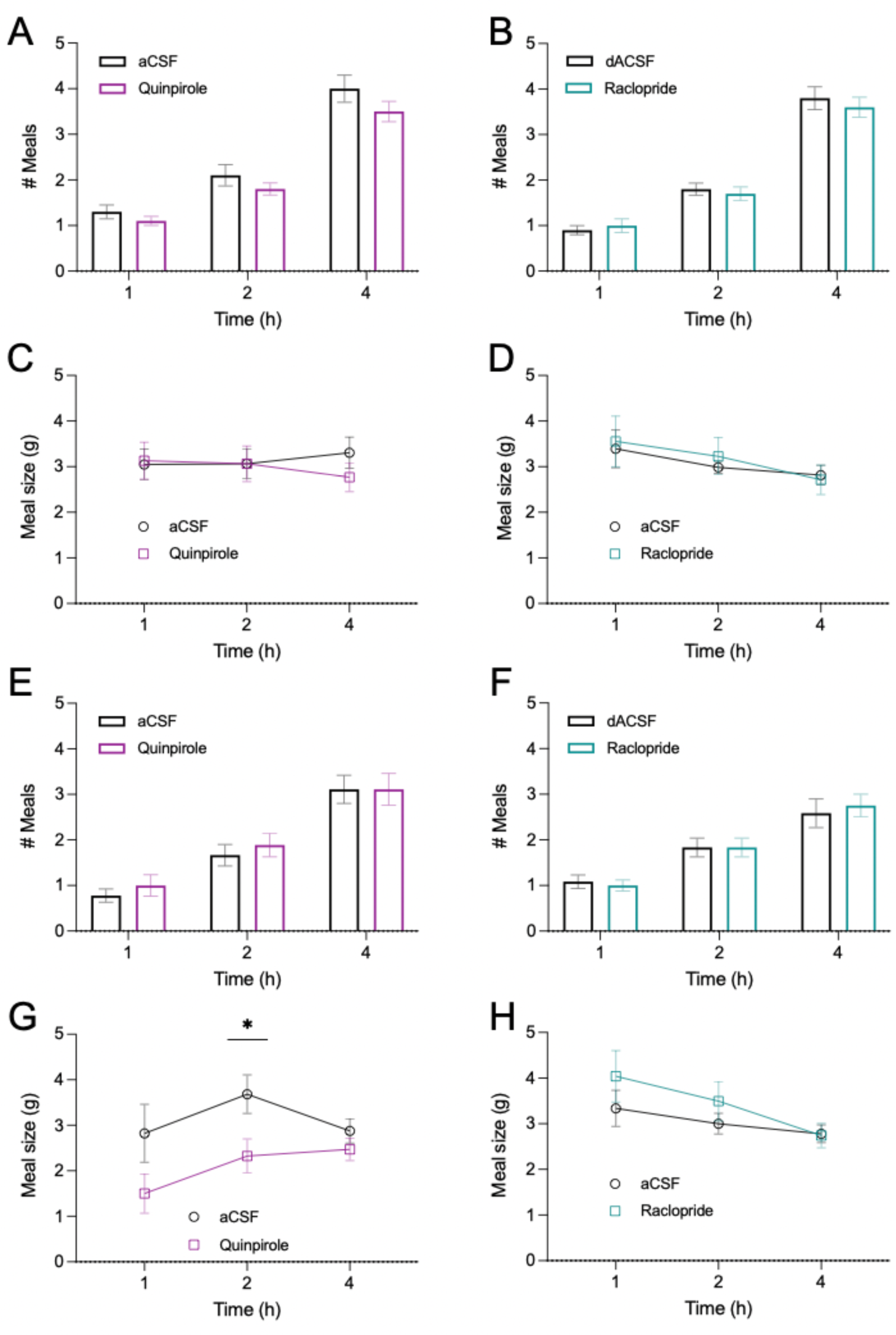
Meal patterns for spontaneous chow intake and post-salient meal chow intake experiments. Cumulative meals consumed in the spontaneous intake experiment did not significantly differ at any time points for **A**) quinpirole vs aCSF [1hr: (t(9)=1.000, p=0.3434), 2hr: (t(9)=1.152, p=0.2789), 4hr: (t(9)=0.6124, p=0.5554)] **B**) raclopride vs aCSF [1hr: (t(9)=1.000, p=0.3434), 2hr: (t(9)=0.5571, p=0.5911), 4hr: (t(9)=0.6124, p=0.5554)]. Cumulative meal size also did not significantly differ at any time points for **C**) quinpirole vs aCSF [1hr: (t(9)=0.1534 p=0.8814), 2hr: (t(9)=0.005189 p=0.9960), 4hr: (t(9)=0.005189 p=0.9960)] or **D**) raclopride vs aCSF [1hr: (t(9)=0.3598, p=0.7273), 2hr: (t(9)=0.5124, p=0.6207), 4hr: (t(9)=0.2574, p=0.8026)]. Post-salient meal cumulative meals consumed did not significantly differ at any time points for **E**) quinpirole vs aCSF [1hr: (t(8)=1.000, p=0.3466), 2hr: (t(9)=0.8000, p=0.4468), 4hr: (t(9)=0.000, p>0.999)]. or **F**) raclopride vs aCSF [1hr: (t(11)=0.3598, p=0.3388), 2hr: (t(11)=0.000, p>0.999), 4hr: (t(11)=0.5606, p=0.5863)]. Spontaneous meal size was significantly reduced at 2 hours for **G**) quinpirole vs aCSF [2hr: (t(8)=2.681, p=0.0279 but did not significantly differ at other timepoints [1hr: (t(8)=2.190, p=0.0600), 4hr: (t(8)=1.115, p=0.2974)], nor at any time points for **H**) raclopride vs aCSF [1hr: (t(11)=1.260, p=0.2339), 2hr: (t(11)=0.8506, p=0.4131), 4hr: (t(11)=0.1182, p=0.9080)]. (All within subject design for meal patterns. Data are means ± SEM; *p<0.05.)

**Supplemental Figure 4:**
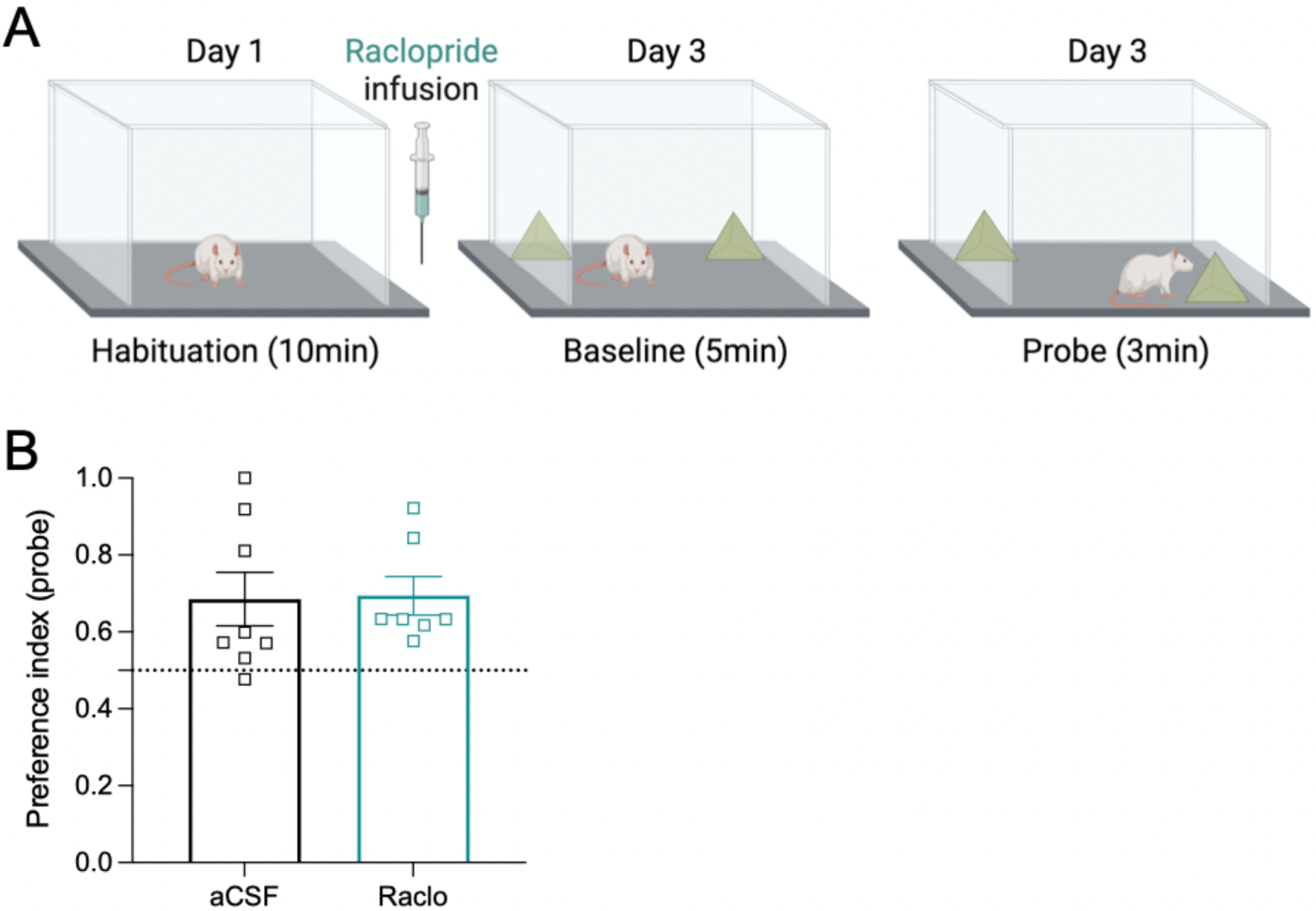
D2R antagonism did not affect short-term spatial object location memory. **A)** Schematic for NLR behavioral experiment. Rats were divided into two groups and treated with raclopride (n=6) or aCSF (n=7) ∼1 hour prior to 5-minute baseline test stage, then returned to their home cage for 5 minutes during which one of the objects encountered at baseline was moved and then returned to the enclosure for a 3-minute probe test stage. **B**) Investigation preference for the displaced object was significantly above the theoretical mean of object investigation at chance (0.5) for both dACSF (t(7)=2.688, p=0.0321) and raclopride treated (t(6)=3.876, p=0.0082) groups. The preference index was not significantly different for raclopride vs aCSF (t(13)=3.876, p=0.0082). (All between-subjects design for NLR. Data are means ± SEM.)

